# PEX1 is essential for the glycosome biogenesis and trypanosomatid parasite survival

**DOI:** 10.1101/2023.08.05.550684

**Authors:** Lavanya Mahadevan, Hemant Arya, Wolfgang Schliebs, Ralf Erdmann, Vishal C. Kalel

## Abstract

Trypanosomatid parasites are kinetoplastid protists that compartmentalize glycolytic enzymes in unique peroxisome-related organelles called glycosomes. The heterohexameric AAA-ATPase complex of PEX1-PEX6 is anchored to the peroxisomal membrane and functions in the export of matrix protein import receptor PEX5 from the peroxisomal membrane. Defects in PEX1, PEX6 or their membrane anchor causes dysfunction of peroxisomal matrix protein import cycle. In this study, we identified the *Trypanosoma* PEX1 orthologue using sequence and structural similarities. Using yeast two-hybrid analysis, we demonstrate that *Tb*PEX1 can bind to *Tb*PEX6. Endogenously tagged *Tb*PEX1 localizes to glycosomes in the *T. brucei* parasites. Depletion of PEX1 gene expression by RNA interference causes lethality to bloodstream form trypanosomes, due to a partial mislocalization of glycosomal enzymes to the cytosol and ATP depletion. *Tb*PEX1 RNAi leads to a selective proteasomal degradation of both matrix protein import receptors *Tb*PEX5 and *Tb*PEX7. Unlike in yeast, PEX1 depletion did not result in an accumulation of ubiquitinated *Tb*PEX5 in trypanosomes. As PEX1 turned out to be essential for trypanosomatid parasites, it could provide a suitable drug target for parasitic diseases. The results also suggests that these parasites possess a highly efficient quality control mechanisms that export the import receptors from glycosomes to the cytosol, in the absence of a functional *Tb*PEX1-*Tb*PEX6 complex.

## 1 Introduction

Trypanosomes are flagellated kinetoplastid protists that cause deadly neglected tropical diseases (NTDs) in vertebrates. Human African trypanosomiasis (HAT), also known as sleeping sickness is caused by *Trypanosoma brucei* and is transmitted to humans by tsetse fly. This disease is endemic in the sub–Saharan African countries (Buscher et al., 2017). Chagas’ disease and leishmaniasis are transmitted to humans by insect vectors called triatomine bugs or sand flies, respectively (Bern, 2015; Burza et al., 2018). Progression of HAT involves two stages where parasites first proliferate in the host bloodstream and lymph (haemolymphatic stage). In the second stage, these parasites penetrate the blood brain barrier (BBB) invading the central nervous system (CNS). Further it leads to an encephalitic reaction (meningoencephalitic stage) that results in sleep disorders, systemic organ failure, and even death if left untreated (Kennedy, 2004). Currently used treatments against HAT include suramin, pentamidine (both administered to cure stage 1 of HAT), Eflornithine, Nifurtimox-Eflornithine combination therapy (NECT) and melarsoprol (to treat the latent second stage). However, the emergence of drug resistance is a major threat and many of these drugs have limitations such as adverse side effects and difficult routes of administration (Fairlamb, 2003; Neau et al., 2020). In the recent years, the development of the first orally administered drug fexinidazole has proven effective to treat the early stages of the disease. However, further clinical trials are ongoing to assess the efficacy of fexinidazole (Sutherland et al., 2017; Dickie et al., 2020). Compared to the HAT where the number of cases has fallen dramatically, Chagas’ disease and leishmaniasis still pose a global public threat. Treatments against these infections also have several drawbacks such as low efficacy against the chronic stages of the Chagas’ disease and emergence of the drug resistant *Leishmania* strains (Wilkinson et al., 2009). These shortcomings imposed by the currently available therapeutic approaches calls for the need to identify novel drug targets that would be more effective and safer for the treatment of trypanosomatid infections (De Rycker et al., 2023).

Trypanosomes contain organelles called glycosomes that are evolutionarily related to peroxisomes (Opperdoes et al., 1984; Quinones et al., 2020). Glycosomes do not possess DNA and rely on the post translational import of the proteins from cytosol for organelle biogenesis. Similar to peroxisomes, glycosome biogenesis also requires peroxins, encoded by PEX genes, which mediate matrix and membrane protein import (Haanstra et al., 2016; Bauer et al., 2017). Enzymes of several metabolic pathways, including the first seven enzymes of glycolysis are compartmentalized within the glycosomes. In the mammalian bloodstream form of *T. brucei* parasites, glycosomes are the sole source of ATP, whereas the mitochondrial activity is repressed. Since parasite glycolytic enzymes lack feedback inhibition, impairment of glycosomal biogenesis leads to their mislocalization to the cytosol, resulting in an accumulation of glucose metabolites at toxic levels, depletion of ATP and cell death (Furuya et al., 2002; Kessler et al., 2005; Haanstra et al., 2008). Thus, targeting glycosomal biogenesis has become an attractive drug target (Kalel et al., 2018). This has been genetically validated using RNA interference mediated knockdown of various peroxins (Moyersoen et al., 2003; Barros-Alvarez et al., 2014; Banerjee et al., 2019), as well as pharmacologically through identification and development of small molecule inhibitors that block peroxin protein-protein interactions (PPIs) (Dawidowski et al., 2017; Banerjee et al., 2021; Napolitano et al., 2022).

In trypanosomes, so far 13 peroxins have been identified and their role in glycosomal protein import pathway has been characterized. Similar to the peroxisomal import pathways in yeast and mammals (Hasan et al., 2013; Fujiki et al., 2014), the cytosolic receptors PEX5 and PEX7 recognize the cargo proteins through their PTS1 and PTS2 signals, respectively, and facilitate the cargo protein import into the glycosomal lumen by shuttling between glycosomal membrane and the cytosol (Moyersoen et al., 2004). After release of cargo, PEX5 is monoubiquitinated, which functions as signal for receptor recycling (Platta et al., 2007; Platta et al., 2009; Gualdron-Lopez et al., 2013; Feng et al., 2022). In yeast and mammals, receptor cycling is mediated by the heterohexameric complex of AAA+ ATPases PEX1 and PEX6 (Miyata et al., 2005; Platta et al., 2005), which is anchored to the membrane by Pex15/PEX26 (Birschmann et al., 2003; Matsumoto et al., 2003). Defects in receptor recycling abrogates peroxisome biogenesis. Particularly, in mammals, mutations in PEX1 (58%), PEX6 (16%) or PEX26 (3%) are together the most prevalent genetic defects that lead to lethal peroxisome biogenesis disorder (PBDs) i.e. Zellweger Syndrome (Ebberink et al., 2011).

Of the three known peroxins involved in the receptor recycling (PEX1, PEX6, PEX15/26), only *Tb*PEX6 has been identified in trypanosomatid parasites (Krazy et al., 2006). The identity of parasite PEX1 and the membrane anchor for the PEX1-PEX6 complex remained unknown. In this study, we report on the identification of trypanosomal PEX1 based on sequence and structural similarity. We show that *Tb*PEX1 localizes to the glycosomes and that it can bind to *Tb*PEX6. RNAi knockdown of *Tb*PEX1 expression leads to parasite cell death by blocking glycosomal protein import and ATP depletion. PEX1 knockdown resulted in a nearly complete degradation of the cargo receptors PEX5 and PEX7, in a proteasome dependent manner. This indicates that trypanosomatid parasites possess a quality control mechanism that dislocates the receptors from the glycosomal membrane when receptor recycling machinery is defective.

## 2 Materials and Methods

### 2.1 Bioinformatic analysis

The plant, insect, yeast and human PEX1 protein sequences were retrieved from UniProt database (UniProt IDs: *Arabidopsis thaliana*, Q9FNP1; Drosophila melanogaster, Q9VUC7; *Saccharomyces cerevisiae*, P24004; *Homo sapiens*, O43933). The TritrypDB database (an integrated platform, which provides access to genome scale datasets for kinetoplastid parasites) (Shanmugasundram et al., 2023) was used to obtain PEX1 homolog sequences of *Trypanosoma brucei* (*Tb*927.4.1250), *Trypanosoma cruzi* (*Tc*C4B63_263g10), and *Leishmania donovani* (*Ld*BPK.34.2.003300). After sequence retrieval, multiple sequence alignment was performed using MEGA v11 MUSCLE tool (Tamura et al., 2021) and the aligned sequences were visualized with Jalview (version 2.11.0) using ClustalX color scheme and conservation threshold of 30%. Phylogenetic comparison between the sequences was also performed using MEGA v11 software. Phobius tool of Stockholm Bioinformatics Centre was used to predict the transmembrane domain and topology. InterPro scan (https://www.ebi.ac.uk/interpro, Release 95.0) was used to identify the domain architecture of the yeast, human and *Trypanosoma* PEX1 sequences (Paysan-Lafosse et al., 2023). PDBsum (http://www.ebi.ac.uk/thornton-srv/databases/pdbsum) and Robetta server (https://robetta.bakerlab.org) were used to predict secondary and tertiary structures, respectively, and the obtained predicted 3D structures were validated using Ramachandran plot analysis (https://saves.mbi.ucla.edu). The validated 3D structures were further visualized and analyzed using PyMOL software (Arya et al., 2014; Schrodinger, 2015).

### 2.2 Yeast Two Hybrid assay

Plasmids expressing *Tb*PEX1 and *Tb*PEX6 fused to GAL4 activation domain (AD) or GAL4 binding domain (BD) were co-transformed into yeast strains PCY2 and PJ694A. PCY2 double transformants were selected on double dropout plates (-Tryp -Leu). Interaction between yeast Pex5p and Pcs60p was used as positive control (Hagen et al., 2015). Yeast 2-hybrid (Y2H) analysis was performed using the protocols described in the Yeast protocols handbook (Clontech, Protocol No. PT3024-1, Version No. PR742227). Positive interaction between two proteins in PCY2 was indicated by the appearance of blue color in the filter based β‐ galactosidase assay, where X-gal was used as a substrate due to its high degree of sensitivity. For the quantitative assessment of the interaction, ONPG was used as the substrate in a liquid assay. The positive interaction in PJ694A was assessed by the growth of double transformants in triple dropout plates (-leu -tryp -his) containing 3 amino 1,2,4 triazole (3-AT), over a period of ten days, where the growth of cells in double dropout plates served as a control. 3-AT, a competitive inhibitor of HIS gene product, was used to avoid false positives in the assay.

### 2.3 Parasite culture and transfection

Bloodstream form (BSF) and Procyclic form (PF) of *T. brucei* Lister 427 strain, cell lines 90.13 and 29.13, respectively, that are genetically modified to express T7 RNA polymerase and Tetracycline repressor, were used in this study. The parasites were grown in HMI-11 (BSF) and SDM79 (PF) medium, containing 10% heat inactivated fetal calf serum (Sigma), 1% Penicillin-streptomycin (Gibco), G418 (15 μg/ml, Invivogen) and Hygromycin (50 μg/ml, Invivogen) respectively. The cultures were maintained in logarithmic phase at 37°C in a humidified incubator with 5% CO_2_ for BSF, or at 27°C for PF. For genetic integration, the constructs were linearized by digestion with NotI, and the DNA was purified by ethanol precipitation. Stable transfection of trypanosomes and clonal selection was performed as described previously (Kalel et al., 2015). The selected clones were stored at −80°C in respective HMI-11 or SDM79 media (without antibiotic) containing 12% glycerol.

### 2.4 Genetic manipulations and plasmid construction

To genomically tag the PEX1 gene in trypanosomes with mNeonGreen, pPOTv7-blast-mNG plasmid was used as a template, with primer pairs RE7224-RE7225 for C-terminal and RE7226-RE7227 for N-terminal tagging. The procedures for the genomic tagging were performed as described by (Dean et al., 2015). For RNA interference, a stem loop construct was generated using two fragments of PEX1 gene. The fragments 1 (1977bp-2407bp) and 2 (1977bp-2458bp) were PCR amplified using primers RE7323-7324 (HindIII and ApaI) and RE7325-RE7326 (ApaI and BamHI), respectively. Following digestion of these two fragments with the above-mentioned restriction enzymes and ligation, the ligated fragment was further digested with HindIII and BamHI and cloned into pHD1336 vector, which contains a tetracycline inducible trypanosome specific promoter. Upon induction of gene expression, a double stranded RNA with a stem loop will be generated, which will cause interference of PEX1 gene expression.

### 2.5 Microscopy

Trypanosomes were harvested by centrifugation and fixed with 4% paraformaldehyde in Phosphate Buffered Saline (PBS, supplemented with 250 mM sucrose in case of RNAi experiments) for 15 min at 4°C. After two washes, the fixed cells were resuspended in PBS and immobilized on poly-L-lysine (Sigma) coated wells, further permeabilized with PBS containing 1% Triton X-100 and blocked with blocking buffer (PBS containing 3% BSA and 0.25% Tween 20). To study the subcellular localization, α-*Tb*Aldolase (1:500 dilution in blocking buffer) was used as glycosomal marker, while Rabbit Alexa Fluor 594 (LI-COR Biosciences) at 1:1000 in blocking buffer was used as secondary antibody. The Nuclear and kinetoplast DNA were stained with DAPI. Stained cells were layered with Mowiol (Sigma) antifade-medium and covered with coverslips. After an overnight setting time that allows the polymerization of Mowiol, the images were captured using Zeiss ELYRA Super Resolution Microscopy and analyzed using Zen 3.6 (blue edition) (Carl Zeiss Microscopy GmbH).

### 2.6 RNA interference and RT-PCR

The PEX1 RNAi construct was stably transfected into bloodstream form trypanosomes and following clonal selection, the cells were seeded at a density of 0.5 million cells/ml and treated with DMSO as negative control (-Tet) or RNAi-induced with 2 μg/ml tetracycline (+Tet) in biological triplicates. The growth of the cells was monitored daily by cell counting using the Neubauer chamber over a period of 6 days, both in the presence (+Tet) and absence of inducer (-Tet i.e., equivalent DMSO). Cells were harvested on days 1 and 2, and RNA was isolated using NucleoSpin® Mini kit (Macherey Nagel). Quantitative Realtime PCR (qRT-PCR) was performed using GoTaq® 1-Step RT-qPCR kit (Promega) with primers specific for PEX1 (RE7039-RE7040) and Tubulin (control, RE7000-RE7001), respectively. qRT-PCR of the samples was performed using Rotor-Gene™ 6000 (Qiagen). The results were analyzed using double delta Ct method (Livak et al., 2001) and graphically plotted using GraphPad Prism 10 software. To study the effect of proteasomal inhibition, cells were treated with 25 μM MG-132 (Sigma) for 6 h, along with tetracycline induction at 2 μg/ml.

### 2.7 Estimation of cell viability and ATP levels

To assess the cell viability, the PEX1 RNAi stably integrated cells were induced as described in Section 2.6. On each day (from day 1 to day 7) post induction, 100 μL of cell suspension was gently transferred to sterile 96-well white opaque bottom plates (Brand GmbH, Germany). To this, 100 μL of CellTiter-Glo® reagent (Promega) was added, and the plate was incubated at room temperature for 20 min. The Luminescence signal was measured using Synergy H1 (BioTek) 96 well-plate reader.

To measure the relative cellular ATP levels in the equal number of parasites, on each day post induction, 0.2 million cells from both DMSO and Tet treatment were harvested, resuspended in 100 μL media, and transferred to a 96-well white opaque bottom plate and the CellTiter-Glo^®^ assay was performed as described above. The luminescence recordings from both cell viability and ATP assessment experiments were analyzed using Graphpad Prism 10 software.

### 2.8 SDS-PAGE and Immunoblotting

The cells harvested from RNAi studies were directly denatured in 1x Laemmli buffer and analyzed by 12% SDS-PAGE. Proteins were transferred to nitrocellulose membranes (Amersham Biosciences) by standard immunoblotting procedure. The membranes were blocked with 5% (w/v) low-fat milk powder to prevent non-specific binding of antibodies and probed with primary antibodies raised in rabbits against purified *Trypanosoma* proteins. To assess the effect of RNAi, α-*Tb*Aldolase, α-*Tb*PFK, α-*Tb*PEX5, α-*Tb*PEX7, α-*Tb*PEX14 and α-*Tb*PEX11 were used to monitor glycosomal markers, while α-*Tb*Enolase was used as cytosolic marker and loading control (each antibody used at 1:10,000 dilution in PBST i.e., PBS containing 1% BSA and 0.05% Tween20). Following the binding of primary antibodies at 4°C overnight under continuous shaking and washing of the membranes, secondary antibody (Alexa Fluor 594, LI-COR, dilution 1:15,000 in PBST without BSA) was added to the membrane for 1 h at room temperature with continuous shaking. Immunodetection of proteins was performed using the Odyssey Infrared Imager and the software Odyssey V3.0 (Li-Cor Biosciences GmbH, Bad Homburg).

### 2.9 Digitonin fractionation

Bloodstream form trypanosomes stably transfected with the PEX1 stem-loop RNAi construct were treated with DMSO or induced for RNAi with 2 μg/ml tetracycline. After 40 h induction, 100 million cells were harvested for both control and RNAi cultures. The harvested cells were subjected to treatment with various amounts of digitonin to assess the release pattern of glycosomal enzymes to the cytosol. After harvesting, the cells were gently washed with ice-cold 1X TNE buffer (50 mM Tris, 150 mM NaCl, 10 mM EDTA; pH 7.4) and briefly centrifuged at 1,500g for 5 min at 4°C. The cell pellets were resuspended in 1x TNE containing EDTA-free Protease inhibitor cocktail (Roche). Following protein estimation using the Bradford method, the cell suspension was aliquoted into several microfuge tubes such that each tube contained cell amount equivalent to 100 μg protein. Various amounts of digitonin (Final concentration of 0-3 mg/mg protein) were added to the aliquots. The tubes were briefly vortexed with mild intensity, and incubated for 3 min at 37°C. 1% Triton X-100 was used as a positive control (complete disruption of cellular membranes and total release of cytosolic as well as glycosomal matrix proteins to the cytosol). The cell suspensions treated with detergents were centrifuged at 13,000 rpm, 10 min at 4°C. The supernatants were carefully aspirated, denatured in 1x Laemmli buffer, and subjected to immunoblotting analysis.

*Primer sequences, strains, plasmids, and cloning strategies are provided in Supplementary tables (See Supplementary material file)*.

## 3 Results

### 3.1 Identification of PEX1 in *T. brucei*

To identify the PEX1 homolog in trypanosomatid parasites, we performed a BLAST search in the TriTrypDB database (www.tritrypdb.com) (Shanmugasundram et al., 2023) against the *Trypanosoma brucei* TREU927 reference proteome, using yeast and human PEX1 protein sequences. Both queries led to the identification of a top hit Tb927.4.1250, which was annotated as a putative peroxisome biogenesis factor 1 and it was further considered as putative *T. brucei* PEX1 (*Tb*PEX1).The second and third top hits were *Tb*VCP/p97 (Roggy et al., 1999) and a nucleolus localized protein (Tryptag.org). The putative *Tb*PEX1 has syntenic orthologs in all *Trypanosoma* and *Leishmania* reference strain species. We retrieved the sequences of the putative *Tb*PEX1 and its counterpart in *T. cruzi* and *L. donovani* from TriTrypDB and aligned them with the known PEX1 protein sequences (**Figure 1A**). Among the PEX1 sequences of all compared organisms, *Tb*PEX1 is the shortest, with 911 amino acids length and a molecular weight of ∼99 kDa. As PEX1 belongs to the Type II of AAA+ ATPases family of proteins, it has two AAA+ domains, namely D1 and D2 positioned after the N-terminus (Grimm et al., 2016). Each domain has a nucleotide binding motif (Walker A) and a nucleotide hydrolysis motif (Walker B), respectively (Grimm et al., 2016). The nucleotide binding domains of *Tb*PEX1 were identified based on the aligned similarity with the yeast and human Walker A motifs (from UniProt database) that are GGSGTGKT (Walker A of domain1, A1) and GASGCGKT (Walker A of domain2, A2), respectively. Similar to other AAA proteins, the second domain (D2) of *Tb*PEX1 appears to be more conserved than the first domain (Grimm et al., 2012) (**Figure 1A**).

**Figure 1.**
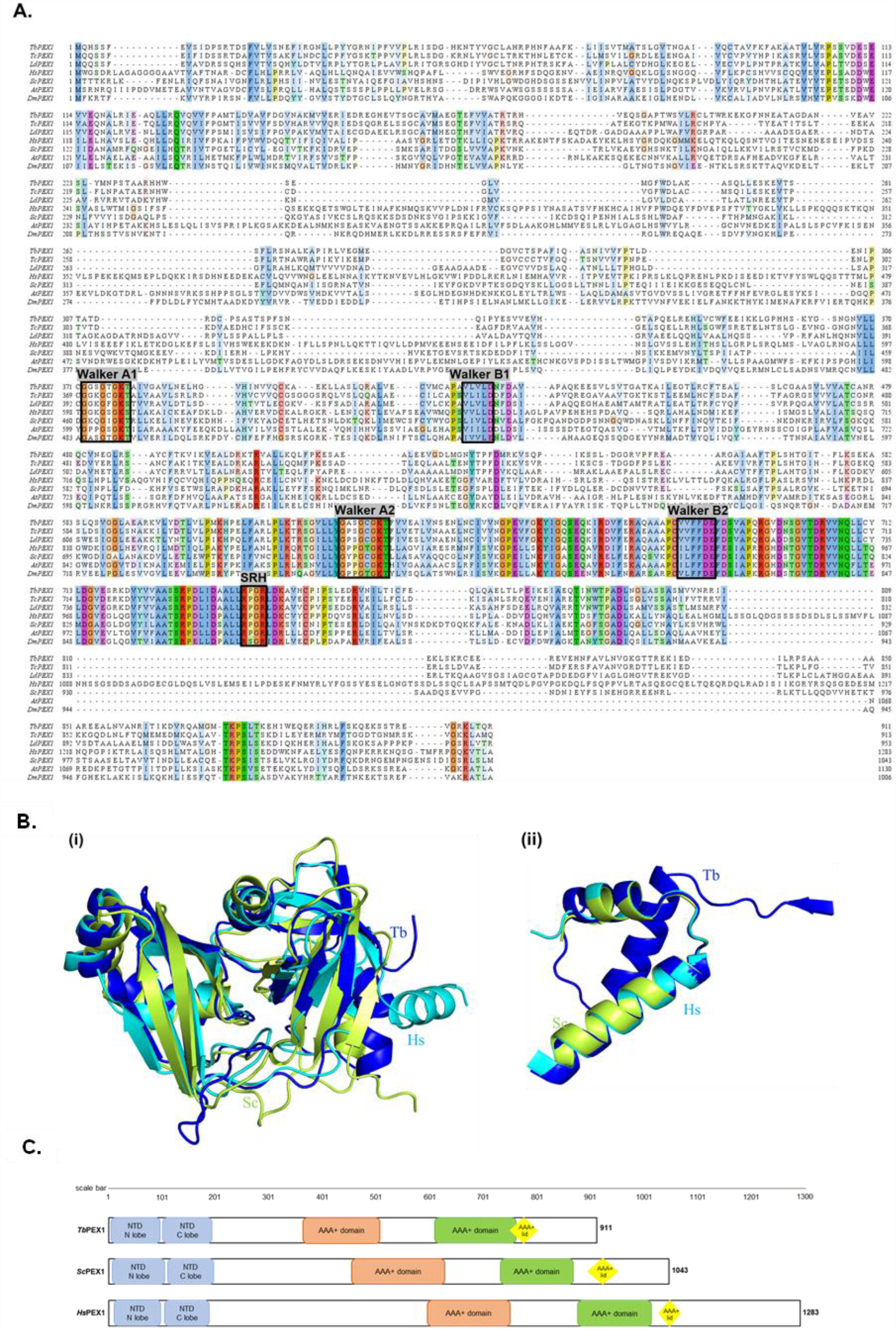
Bioinformatic analysis of PEX1 orthologs of yeast, human, plant, insect, and parasites. A. Multiple sequence alignment of yeast (*S. cerevisiae*), human (*H. sapiens*), plant (*Arabidopsis thaliana*), insect (*Drosophila melanogaster*) and trypanosomatid parasite protein sequences. Sequences of yeast, human, plant, and insect PEX1 were obtained from UniProt (UniProt IDs P24004, O43933, Q9FNP1 and Q9VUC7), parasite protein sequences were obtained from TriTrypDB with Accession IDs Tb927.4.1250 (*T. brucei*), TcCLB.505989.74 (*T. cruzi*) and LdBPK_343300.1 (*L. donovani*). Multiple Sequence alignment was performed using MUSCLE tool and the aligned sequences were visualized using Jalview software with ClustalX color scheme. The nucleotide binding regions (Walker A) and nucleotide hydrolysis regions (Walker B) of domains 1 and 2 are indicated in black and red boxes, with the nomenclature WA1/WB1 and WA2/WB2, respectively. Second region of homology (SRH) is only present in D2, indicating that like other organisms, also parasite D1 of PEX1 can bind ATP but not hydrolyze it (Blok et al., 2015) **B i)** Comparison of the three-dimensional (3D) structure of the N-terminal region (NTD) of the putative PEX1 of *Trypanosoma brucei* (blue) with mammalian PEX1 (cyan) (obtained from the comparative modeling), and *Saccharomyces cerevisiae* PEX1 (lemon). **ii)** 3D structure of the identified AAA+ lid of the putative PEX1 of *Trypanosoma brucei* (blue) with mammalian PEX1 (cyan), and PEX1 from *Saccharomyces cerevisiae* (olive green) **c)** Schematic representation of the domain architecture of the putative *Tb*PEX1, *Hs*PEX1 and *Sc*PEX1. The NTD (*Tb*PEX1: 6-94aa, 101-180; *Hs*PEX1: 17-98aa, 104-179aa; *Sc*PEX1-12-102aa, 108-183aa) is highly conserved and is structurally similar across the organisms. The RMSD score between the NTD structures is less than 2 Å. In additional to AAA+ domains, the AAA+ lid (yellow) is also well conserved and is found to be structurally identical in all the three organisms.

According to the results from Sequence Identities and Similarities (SIAS) webtool, PEX1 of parasites is poorly conserved: the parasite protein sequences (*Tb, Tc, Ld*) themselves share only overall 9-17% identity, and a mere ∼6% identity with the entire length of yeast and human PEX1 proteins (**Supplementary Figure 1A**). The phylogenetic analysis suggests that the putative TbPEX1 is evolutionarily closely related to its identified homologues in *Tc, Ld*, and *At* and less related to *Dm, Sc*, and *Hs*PEX1 (**Supplementary Figure 1B**). The prediction of combined transmembrane topology and signal peptide using Phobius software indicated that *Tb*PEX1 does not possess transmembrane domains (not shown), which is similar to the homologs in plant, yeast and mammals (Tamura et al., 1998; Birschmann et al., 2005). The PEX1 protein 2D structures of *Tb, Hs*, and *Sc* predicted using PDBsum suggested that they share fewer similarities (**Supplementary Figure 2A**).

**Figure 2.**
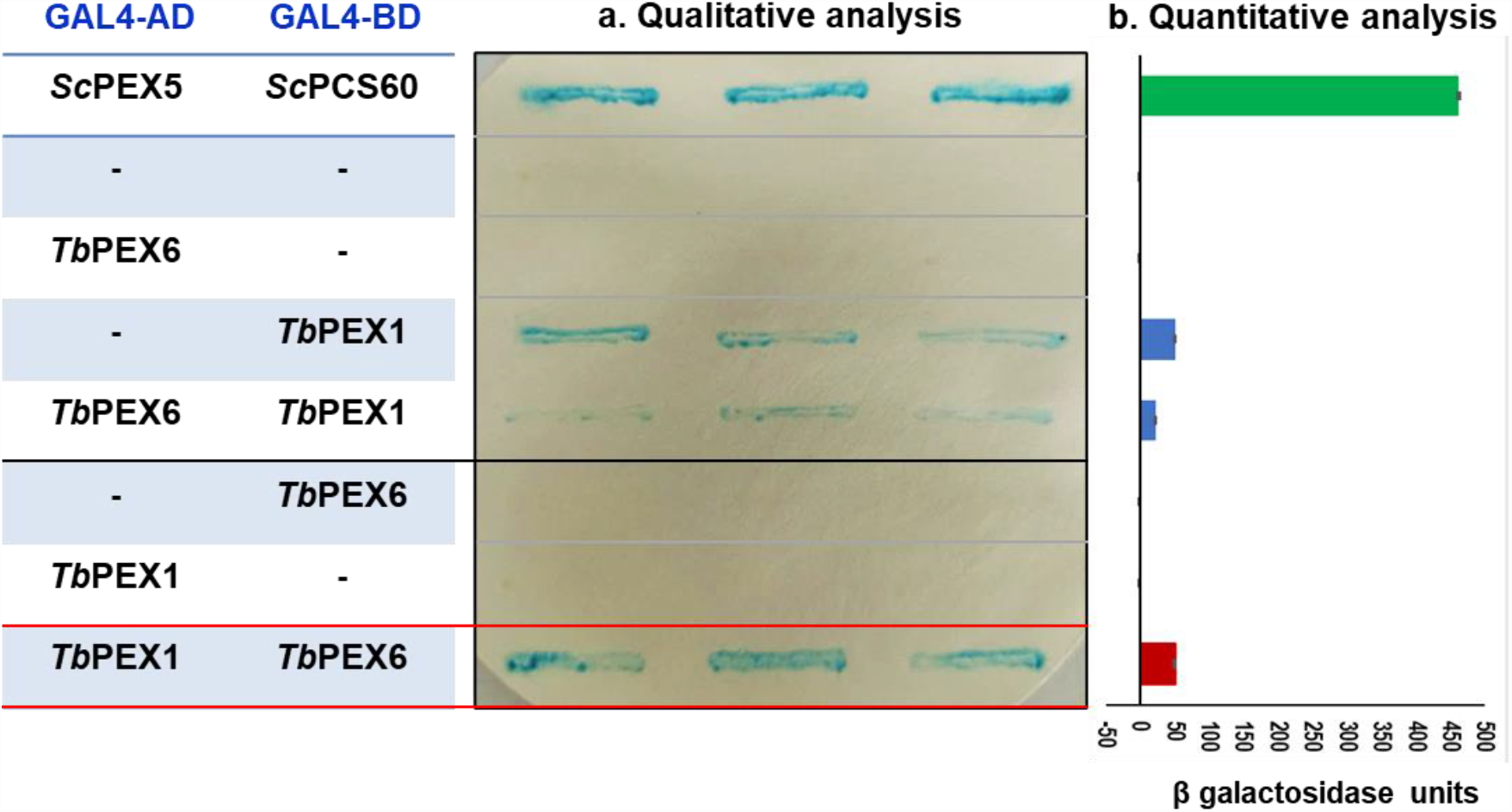
*Tb*PEX1 interacts with *Tb*PEX6 in yeast two-hybrid (Y2H) system. *Tb*PEX1 or PEX6 were fused with GAL4-AD or -BD in Y2H vectors pPC86 or pPC97. PCY2 yeast strain was co-transformed with the construct or empty vector as indicated, and double transformants were selected using double dropout media (-Leu -Tryp). Three independent PCY2 transformed clones were analyzed by a qualitative colony-lift filter assay using X-Gal as substrate (**A)** or a quantitative liquid assay using ONPG as the substrate which was then quantified spectrophotometrically **(B)**. *Tb*PEX1 (AD fusion) showed interaction with *Tb*PEX6 (BD fusion) in both solid and liquid assays and respective controls (interaction with empty vector) did not show any autoactivation. The interaction between *Sc*PEX5 and its cargo *Sc*PCS60 served as a positive control. Error bars shown in the graph represent the Standard deviation between three biological replicates used in the assay.

Since the overall sequence similarity of PEX1 proteins was low, we further assessed the structural homology of *Tb*PEX1 with the mammalian counterpart. Molecular modeling is a theoretical-based computational technique to generate or derive the three-dimensional structure of target proteins using machine learning, genetic algorithms, and artificial intelligence. In this study, the Robetta server predicted the full-length 3D structure of *Tb*PEX1, *Tc*PEX1, and *Hs*PEX1 with a high confidence score (more than 0.6). Moreover, the predicted structures were minimized using the maestro tool of Schrödinger software and validated through the Ramachandran plot. Modeled structures showed that more than 98.5% of residues were present in the allowed region of the Ramachandran plot (**Supplementary Figure 2B**).

The crystal structure of the mammalian PEX1 N-terminal domain (NTD) (PDB ID-1WLF), with a few missing residues, is available in the Protein Data Bank (Shizowa et al. 2004). The NTD structure was retrieved, and the missing amino acids were filled in by comparative modeling using the Robetta server. The crystal structure and modeled protein nicely superimposed on each other with 0.29 Å RMSD (**Supplementary Figure 2C**). Moreover, the NTDs of *Tb*PEX1 and *Sc*PEX1 superimposed with the mammalian (*Hs*PEX1) NTD structure. The structure of the NTD region of *Tb, Hs*, and *Sc* were found to be structurally similar with an RMSD of 1.69 Å (between *Hs* and *Tb*) and 1.86 Å (between *Hs* and *Sc*) (**Figure 1B, left panel**). The literature survey and InterPro scan indicated the presence of three domains in *Hs* and *Sc*, whereas *Tb* consists only of the NTD and AAA+ domains (**Figure 1C**). We further performed extensive structural analysis and identified the AAA+ lid feature in *Tb*PEX1, which was not detected in the InterPro Scan (**Figure 1B, right panel and 1C**). The AAA+ lid in *Tb*PEX1 encompasses 747 to 807 residues, and it is structurally similar to the AAA+ lid of *Sc*PEX1 and *Hs*PEX1 (**Figure 1C**). Furthermore, the high throughput endogenous tagging studies indicate that this protein localizes as puncta in parasites (Tryptag.org), and it was also identified in the proteomic analysis of glycosomes (Guther et al., 2014). This information along with the characterization described further in this study demonstrate that the identified putative protein is the true *Tb*PEX1 ortholog.

### 3.2 *Tb*PEX1 interacts with *Tb*PEX6 in a yeast two-hybrid (Y2H) assay

PEX1 is known to bind to PEX6 and to form heterohexameric complexes in yeast and mammalian cells. To investigate if this interaction is conserved in *Trypanosoma* parasites, we performed a Y2H analysis with full length *Tb*PEX1 and *Tb*PEX6 proteins fused to either GAL4-AD or -BD. Double-transformed clones of the PCY2 yeast strain were selected on -leu -tryp dropout media and were assayed for β-galactosidase activity using a colony-lift filter assay with X-gal as the substrate (**Figure 2A**). An autoactivation was observed with *Tb*PEX1 fused to GAL4-BD, therefore this plasmid is not suitable to study the interaction with *Tb*PEX6 fused to GAL4-AD. From the blue color appearance of colonies in the plate assay, it is clear that *Tb*PEX6 fused to GAL4-BD interacts with *Tb*PEX1 fused to GAL4-AD. The positive interaction was further assessed quantitatively in a liquid assay using ONPG as a substrate (**Figure 2B**), which confirmed the results of the plate assay. However, the interaction of TbPEX1 fused to GAL4-AD with *Tb*PEX6 fused to GAL4-BD is weaker in comparison to the positive control, the interaction of *Sc*PEX5 and *Sc*PCS60. This could be due to a lower affinity of the protein pair but could also be caused by a poor nuclear translocation of the large PEX1-PEX6 complex as compared to PEX5-PCS60, or differences in their expression levels. However, the interaction between *Tb*PEX1-GAL4-AD and *Tb*PEX6-GAL4-BD was also confirmed in the PJ694A strain, where the interaction is indicated by the growth of cells in triple dropout plates (-leu -tryp -his) (**Supplementary Figure 3**).

**Figure 3.**
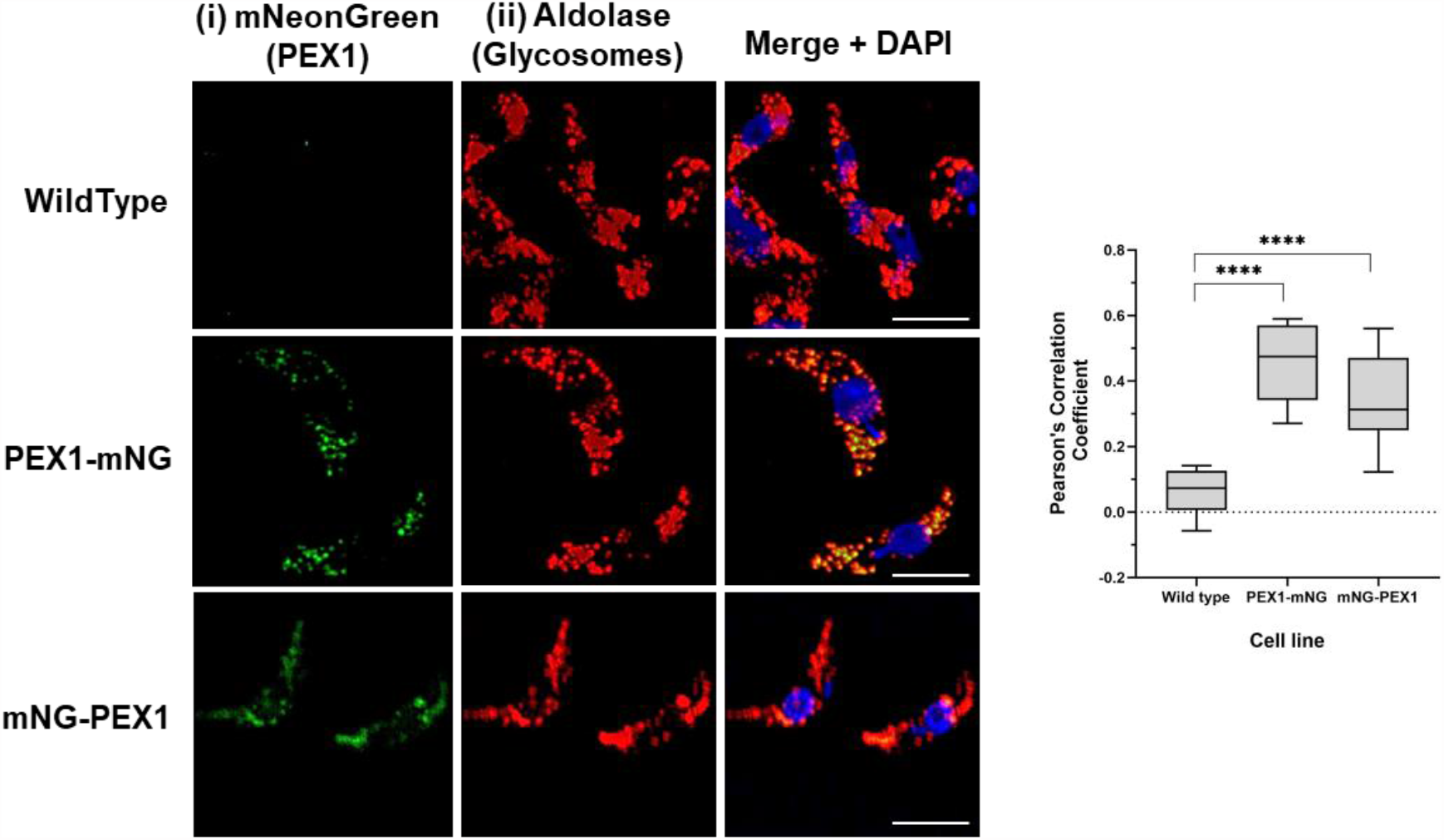
*Tb*PEX1 endogenously tagged with mNeonGreen localizes to glycosomes in procyclic form trypanosomes. Immunofluorescence microscopy staining of glycosomes was performed using antibodies against glycosomal enzyme *Tb*Aldolase, which gives a typical punctate pattern of glycosomes. Both N- and C-terminally mNG tagged *Tb*PEX1 (PEX1-mNG, mNG-PEX1) colocalized with the glycosomal marker as seen in merge panel. DAPI stains both nuclear and kinetoplast DNA. Scale bars are 2 μm. The data graphically presented (right panel) using GraphPad Prism 10.0 software represents the Pearson colocalization coefficient calculated using the colocalization tool of the Zen Blue software version 3.6. The line in the box and whiskers plot denotes the median value of each dataset (n =10 cells). One-way ANOVA (Dunnett’s test) performed with the datasets gave a p<0.0001 at 95% confidence indicated by ****.

### 3.3 Subcellular localization of *Tb*PEX1 to glycosomes

To monitor the sub-cellular localization of the *Tb*PEX1 protein in parasites, the corresponding PEX1 gene was endogenously C-terminally or N-terminally tagged with the mNeonGreen (PEX1-mNG, mNG-PEX1). Following stable genomic integration in procyclic form trypanosomes, single clones were selected by limiting dilution of the transformants. The clones selected were confirmed for endogenous tagging by PCR (**Supplementary Figure 4**). The procyclic form trypanosomes encoding the endogenously tagged PEX1 were fixed with formaldehyde and visualized for colocalization of *Tb*PEX1 with the glycosomal marker aldolase by immunofluorescence microscopy. Aldolase labelling of the cells resulted in a punctate pattern (**Figure 3 panel ii**, red labelling), which is a typical morphological hallmark feature of peroxisomes/glycosomes. mNeonGreen fluorescence indicated the localization of the tagged *Tb*PEX1 in cells (**Figure 3 panel i**, labelled in green). The colocalization of both N-or C-terminally mNG-tagged PEX1 with the punctate labelling of aldolase indicates that *Tb*PEX1 is a glycosomal protein, as evident in the merge channel (**Figure 3** middle and lower panel). This is also consistent with the finding that the *Tb*PEX1 is significantly detected in the high confidence glycosomal proteome of trypanosomes (Guther et al., 2014).

**Figure 4.**
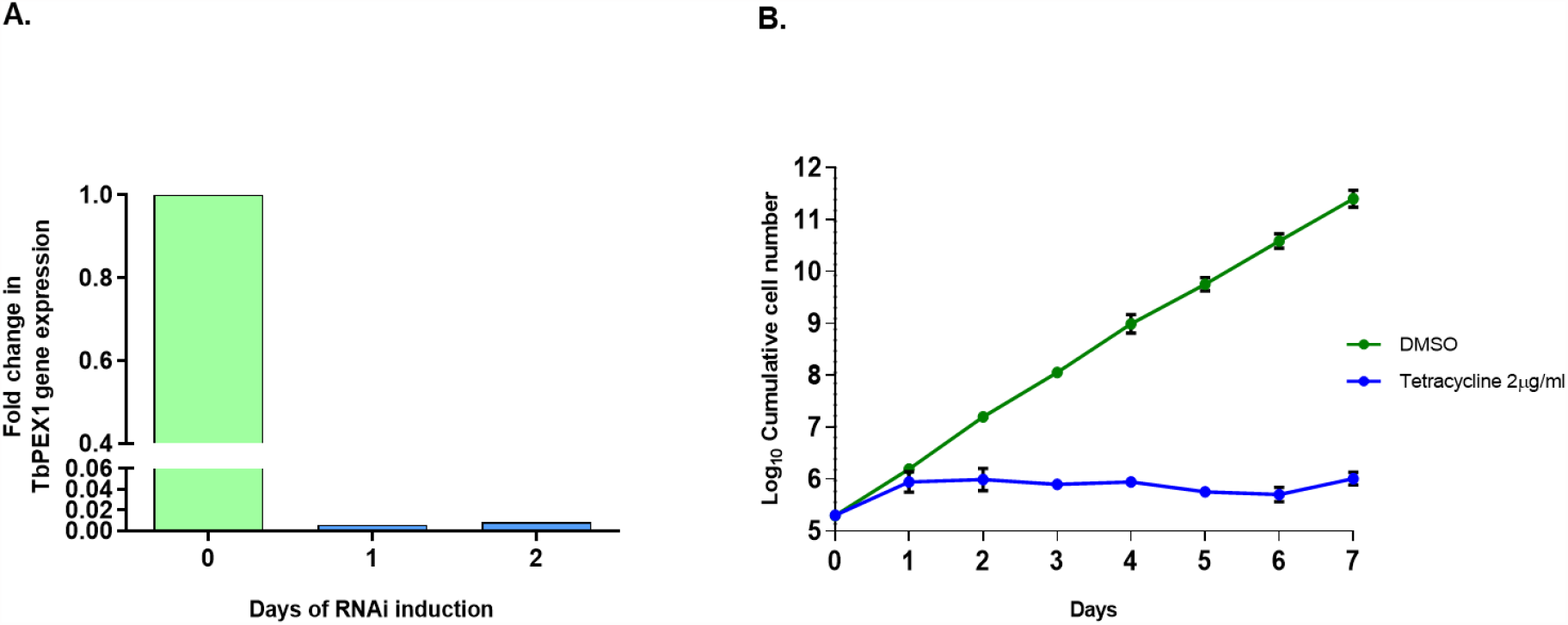
*Tb*PEX1 is essential for the survival of bloodstream form trypanosomes A. Quantitative RT-PCR analysis to determine the expression of *Tb*PEX1 mRNA in cells induced for RNAi by tetracycline addition on days 1 and 2, respectively. The data shown are normalized values with control (DMSO) from respective days. qRT-PCR with Tubulin specific primers served as internal control to calculate double delta CT values which allow normalization between DMSO and Tet induced RNAi samples. **B**. BSF trypanosomes stably transfected with tetracycline inducible *Tb*PEX1 RNAi construct was induced with tetracycline or treated with DMSO as control. Cells were manually counted every day and diluted back to the seeding density of 0.2 million cells/ml. Cumulative growth was plotted in Log_10_ scale using GraphPad Prism (version 10). The cell survival analysis demonstrates that *Tb*PEX1 RNAi led to a severe growth defect in BSF parasites. The error bars represent the standard deviation among 3 biological replicates.

### 3.4 RNA interference of *Tb*PEX1 leads to severe growth defect in bloodstream form trypanosomes

To evaluate the effects of PEX1 gene knockdown on the parasites, bloodstream form trypanosomes were transfected with a tetracycline inducible stem-loop RNAi construct. Following the clonal selection, RNA interference was induced by adding 2 μg/ml of tetracycline. DMSO treatment (non-induced) served as control in this experiment. After induction of RNAi, the growth of the cells was assessed over a period of 6 days. The cell viability was manually determined on each day of the experiment by counting the cells using a Neubauer chamber. The initial seeding density of cells was 0.2 million cells/ml and after every day’s count, the cells were adjusted to the initial seeding density. To assess the *Tb*PEX1 gene expression levels of the RNAi cells, they were harvested from days 0 to 2 of the study. After total RNA isolation from the cells, quantitative RT PCR was performed using primers specific to *Tb*PEX1. Primers specific to Tubulin served as a control. The qRT-PCR data graphically plotted using GraphPad Prism (**Figure 4A**) demonstrates that an RNAi efficiency as close as 100% has been achieved in reducing the PEX1 gene expression post RNAi induction with tetracycline. Cumulative growth of the cultures was plotted to assess cell survival (**Figure 4B**). The cells induced for *Tb*PEX1 RNAi displayed decline in growth pattern from day 1 onwards and this effect was observed until day 6, while the cells in the control group (DMSO treated *Tb*PEX1 RNAi cell line i.e., uninduced) continued to grow exponentially. The cell viability, assessed using the Luminescent CellTiter-Glo^®^ based method, also indicated the significantly reduced cell number over time upon PEX1 RNAi induction (**Supplementary Figure 5**).

**Figure 5.**
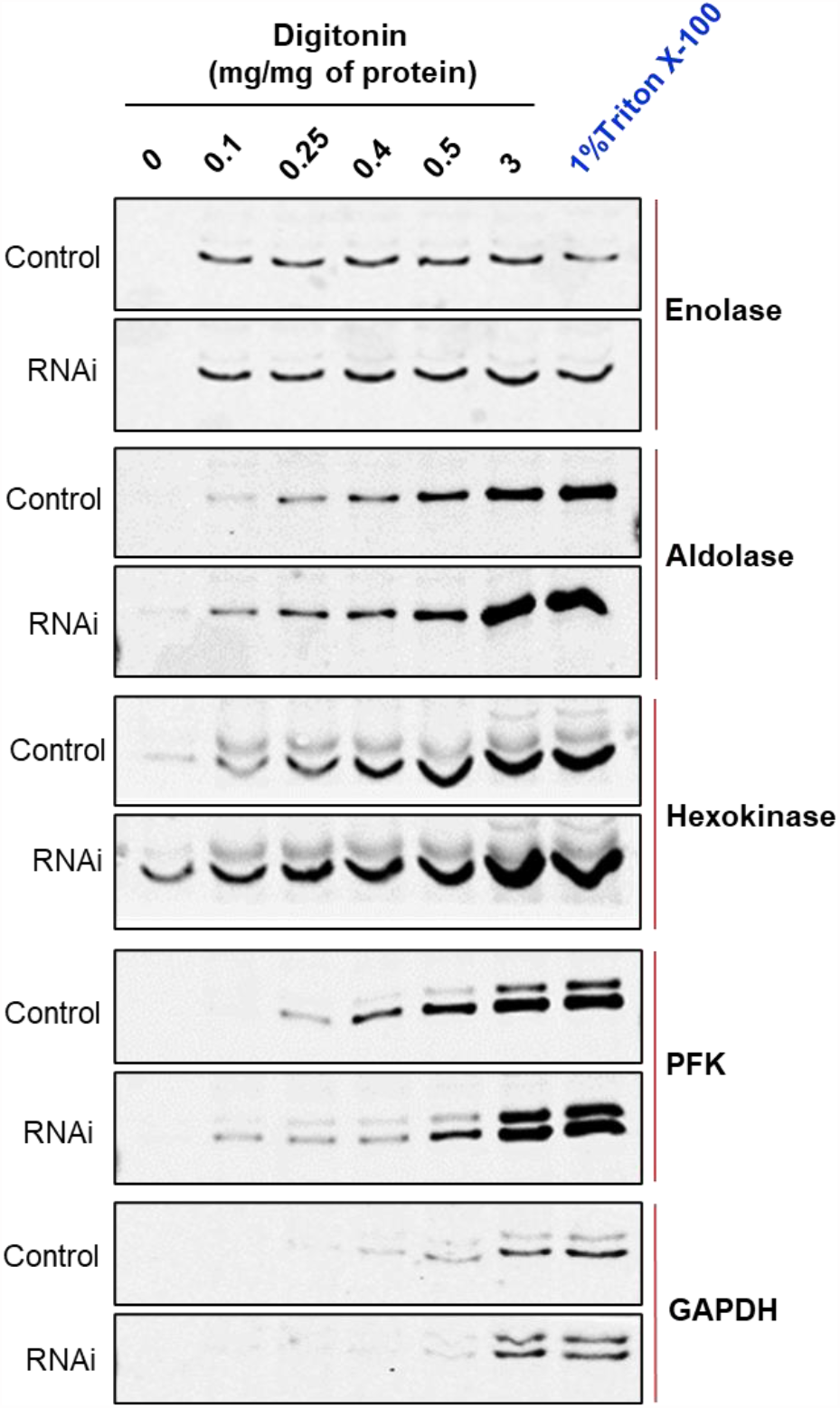
RNAi knockdown of *Tb*PEX1 leads to a partial mislocalization of glycosomal enzymes to the cytosol. Immunoblot analysis showing the differential release pattern of glycosomal enzymes from bloodstream form PEX1 RNAi cell line that were treated with DMSO (Control) or tetracycline to induce RNAi (RNAi) for 2 days. Digitonin fractionation was performed with indicated amounts of digitonin, which permeabilizes the plasma membrane and releases cytosol already at low concentrations. At higher concentrations organellar membranes are permeabilized and luminal proteins of the organelles are released. No detergent served as negative control (0 mg digitonin/mg of protein), while 1% Triton X-100 treatment as positive control for complete release of cytosolic and matrix proteins. Supernatants of the treated samples were analyzed by immunoblotting with antibodies against cytosolic marker Enolase (which also served as loading control) and indicated glycosomal matrix markers: PFK-Phosphofructokinase, GAPDH – Glyceraldehyde 3 phosphate dehydrogenase (both PTS1 proteins); Aldolase and Hexokinase (both PTS2 proteins). Even low concentration of digitonin i.e., 0.1 mg/mg led to complete release of cytosol in both ‘Control’ and ‘RNAi’ cells. Glycosomal enzymes Aldolase, Hexokinase and PFK were detected in the supernatants of RNAi cells even at lowest digitonin concentration, indicating that these three enzymes are partially mislocalized to the cytosol upon PEX1 RNAi. GAPDH did not show this behavior. Shown here is a representative of the results obtained in three independent biological replicates.

### 3.5 Knockdown of PEX1 results in partial mislocalization of glycosomal enzymes and ATP depletion

Defects in PEX1, PEX6 or their anchor protein are known to disrupt peroxisomal protein import in other organisms (Reuber et al., 1997; Mastalski et al., 2020). To biochemically assess whether the RNAi knockdown of PEX1 led to a mislocalization of glycosomal enzymes, the RNAi cell line treated with DMSO alone i.e., non-induced and Tet-induced cells (labelled as ‘Ctrl’ and ‘RNAi’, respectively in **Figure 5**) were both analyzed by digitonin fractionation. Equal amounts of cells were aliquoted in different microfuge tubes and treated with increasing amounts of the digitonin, that allows release of cytosolic and organellar matrix proteins in an increasing concentration dependent manner. Treatment with 1% Triton X-100 ensures complete dissolution of cellular membranes, while treatment without digitonin served as negative control. After detergent or buffer alone treatment, cell suspensions were centrifuged, and the supernatants were analyzed by immunoblotting. Enolase is a cytosolic marker and is released completely in both control and RNAi induced samples even at lowest concentration of digitonin i.e., 0.1 mg/mg protein (**Figure 5**), except for the negative control without digitonin treatment where the plasma membrane remains intact. Whereas glycosomal enzymes require more than 0.5 mg digitonin per mg cells for complete release in control. In RNAi induced samples, glycosomal enzymes aldolase, hexokinase (both PTS2 signal containing enzymes) and PFK (PTS1 containing enzyme) were specifically released in higher amounts and at lower concentrations of digitonin than in the non-treated control cells. This indicates that glycosomal enzymes are mislocalized to the cytosol upon PEX1 RNAi. However, the PTS1 containing enzyme glycosomal GAPDH was not affected by the RNAi. Here, we must consider that the growth of the cells is severely affected after two days of RNAi, which means that the glycosomes of the cells still contain proteins that were imported prior to the RNAi treatment. Accordingly, only newly synthesized proteins will remain in the cytosol upon RNAi. This consideration explains that the glycosomal proteins in this experiment show a bipartite behavior upon RNAi. Accordingly, the newly synthesized proteins remain in the cytosol and are released at low digitonin concentrations, while the glycosomal proteins that were imported earlier are released at higher concentrations.

If glycosomal enzymes are mislocalized to the cytosol, their unregulated activity results in ATP depletion, which kills trypanosomes (Haanstra et al., 2008). *Tb*PEX1 RNAi led to a partial mislocalization of glycolytic enzymes to the cytosol, including the ATP consuming kinases, Hexokinase and PFK. Therefore, we measured the total cellular ATP levels in cells induced for PEX1 RNAi relative to DMSO treated control cells (**Figure 6**). We observed a significant reduction in ATP levels from day 1 to day 3 of *Tb*PEX1 RNAi induction. This confirms the mislocalization of glycolytic enzymes upon *Tb*PEX1 RNAi and shows that even a partial mislocalization of glycolytic enzymes (as seen in **Figure 5**) results in ATP depletion and killing of *Trypanosoma* parasites.

**Figure 6.**
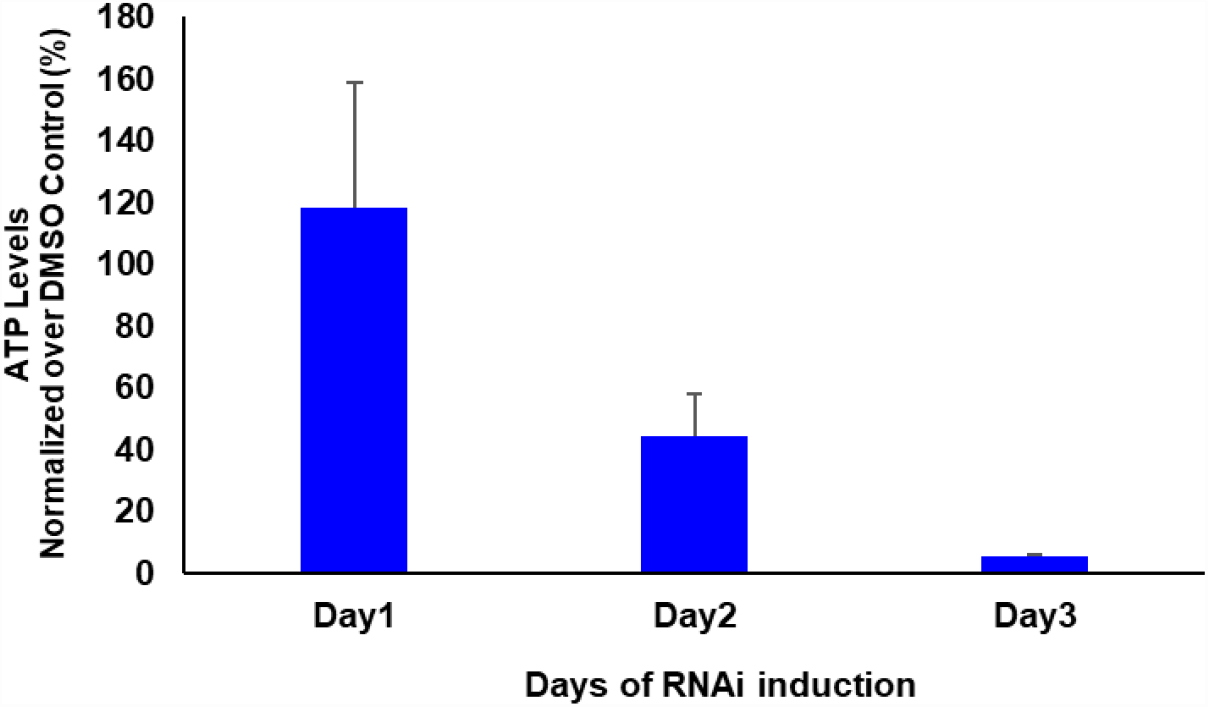
RNAi knockdown of PEX1 led to ATP depletion in *Trypanosoma* parasites. Bloodstream form *T. brucei* PEX1 RNAi cells were treated with DMSO as control or tetracycline to induce PEX1 RNAi. Equal number of control and RNAi cells were harvested, and the ATP content was estimated using CellTiter-Glo® reagent (Promega). The luminescence readings were normalized and the percent relative ATP levels in RNAi induced cells were graphically represented using GraphPad Prism version 10. A clear reduction in ATP levels was observed upon PEX1 RNAi depletion, on days 2 and 3. The error bars indicate the standard deviation among the three biological replicates that were used in the assay.

### 3.5 Silencing of PEX1 expression results in degradation of the cargo receptors in a proteasome dependent manner

To study the effects of PEX1 RNAi knockdown on the receptor recycling process, control and RNAi induced cells were harvested on days 1 and 2. Total cell lysates were analyzed for the steady state levels of the glycosomal cargo receptors PEX5 and PEX7. Docking factor PEX14 and PEX11 were analyzed as glycosomal membrane markers, PFK and Aldolase were analyzed as glycosomal enzyme, while the cytosolic marker Enolase served as a loading control (**Figure 7A**). A significant reduction in the steady state levels of PEX5 was already evident on day 1 of RNAi induction (+Tet), while on later time points PEX5 could be hardly detected in the immunoblots (**Figure 7A, Top panel**). Similar trend of reduction and disappearance of PEX7 is seen, albeit in a delayed manner on day 2 and 3, as compared to PEX5. In contrast, glycosomal membrane proteins PEX14 or PEX11 were unaffected, and the glycosomal matrix enzymes PFK and Aldolase were either not affected or mildly affected, indicating that the observed decrease in PEX5 and PEX7 is not caused by pexophagy. This shows that both cargo receptors PEX5 and PEX7 are specifically degraded, when PEX1 expression is knocked down.

**Figure 7.**
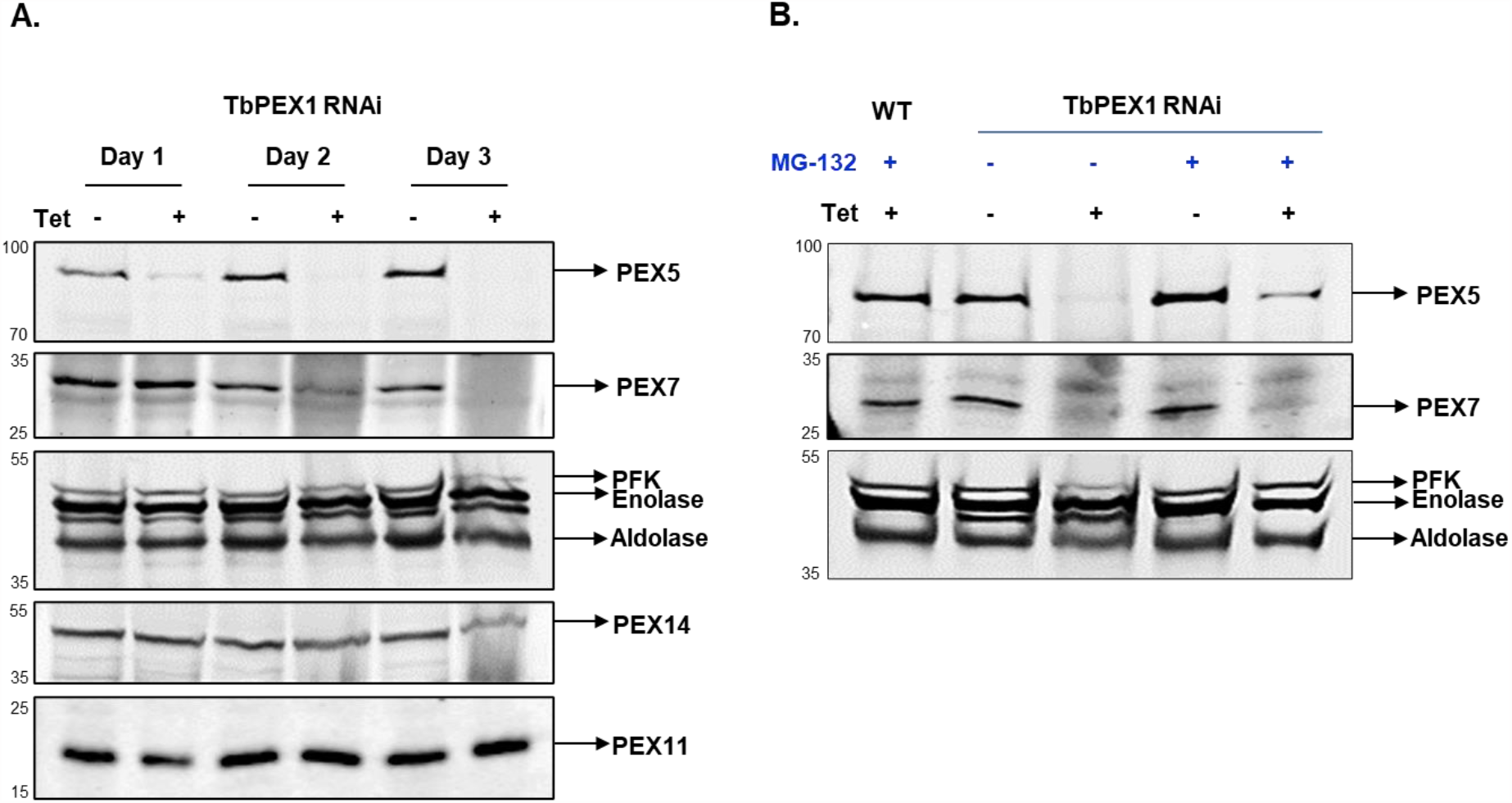
Glycosomal import receptors are degraded upon PEX1 RNAi A. Immunoblot analysis of total cell lysates from PEX1 RNAi cells treated with DMSO (Tet-) or RNAi-induced with tetracycline (Tet+) was performed using antibodies as indicated. It is evident that the cargo receptors PEX5 and PEX7, that recognize PTS1 and PTS2, respectively, are unstable upon induction of PEX1 RNAi. Significant degradation of PEX5 is evident on day 1 itself, and for PEX7 on day 2 onwards. Enolase served as a loading control. Other glycosomal markers such as PFK, Aldolase and membrane proteins PEX14 and PEX11 are also assessed, which show no or minor degradation on day 3. This suggests that only the cargo receptors are specifically degraded, with no or minimal contribution from pexophagy **B**. PEX1 RNAi cell line was grown for 2 days (Tet-/+), and subsequently treated with proteasome inhibitor MG-132 or DMSO for 6 h. Total cell lysates were analyzed by immunoblotting. The analysis shows that the cargo receptors PEX5 and PEX7 are partially protected from proteasomal degradation in PEX1 knockdown cells by MG-132 treatment (last lane).

In the absence of PEX1, PEX5 should remain stuck at the membrane, however specific degradation of the cycling import receptors suggests that they are dislocated out of the membrane and further degraded. To investigate whether degradation of the proteins involves the proteasome, MG-132, a specific inhibitor of 26S proteasome was used. After 6 h treatment with 25 μM MG-132, PEX1 RNAi cell line (with and without tetracycline addition) were harvested and assessed for PEX5 and PEX7 levels. As shown in **Figure 7B**, PEX5 is rescued from degradation in RNAi induced cells when the proteasomal activity is inhibited by MG-132. Since the parasites can only be treated with MG-132 for limited time, full rescue of PEX5 cannot be expected, as longer MG-132 treatment itself would lead to cell toxicity. Taken together, the data show that the import receptor PEX5 is degraded in PEX1-deficient cells in a proteasome dependent manner.

## 4 Discussion

In this work, we report the identification and characterization of the trypanosomal PEX1, a key component of the glycosomal protein import and quality control machinery. Over the past years, using the sequences of known peroxins from yeast, mammals and plants, primary sequence-based BLAST search led to the *in silico* identification of several trypanosomatid peroxins. Similarly, we identified trypanosomal PEX1 by BLAST search using the sequences of yeast and mammalian orthologs. Despite the low overall sequence similarity, the AAA+ domain, in particular the D2 ring shows higher degree of sequence conservation. Further, structural modelling showed that the domain architecture and 3D structures of PEX1 are highly conserved across different organisms (Shiozawa et al., 2004). Similar approaches were used for the identification of *Tb*PEX3, using remote homology (HHPRed) and structural similarity (Phyre2) (Banerjee et al., 2019; Kalel et al., 2019). We further showed that *Tb*PEX1 interacts with *Tb*PEX6, and endogenously mNeonGreen-tagged PEX1 localizes to glycosomes in *Trypanosoma* parasites.

Glycosomes are essential for the parasite survival. Accordingly, disruption of glycosome biogenesis by RNAi knockdown or chemical inhibition of PEX proteins kills *Trypanosoma* parasites (Haanstra et al., 2016; Dawidowski et al., 2017). RNAi knockdown of PEX1 expression led to a severe growth defect in bloodstream form trypanosomes, which validates that PEX1 is essential for the survival of the parasites. PEX1 RNAi resulted in a partial glycosomal protein import defect, which is enough to kill the bloodstream form *Trypanosoma* parasites, as they solely rely only on glycosomes for the energy production. A similar growth phenotype and glycosomal protein import defect was also in seen trypanosomes upon RNAi knockdown of the partner AAA+ ATPase of *Tb*PEX1 i.e. *Tb*PEX6 (Krazy et al., 2006). Mislocalized glycosomal enzymes exhibit uncontrolled activities in the cytosol, thus leading to ATP depletion and accumulation of glucose metabolites to toxic levels, which causes death of parasites (Furuya et al., 2002; Kessler et al., 2005; Haanstra et al., 2008). Accordingly, we also observed ATP depletion upon PEX1 RNAi knockdown.

*Tb*PEX1 RNAi led to cytosolic mislocalization of both PTS1 and PTS2 containing glycosomal enzymes and to a selective degradation of both the PTS1- and the PTS2-import receptor, *Tb*PEX5 and *Tb*PEX7, respectively. Ubiquitination of PEX5 and its role in receptor recycling (mono-ubiquitination) or proteasomal degradation (poly-ubiquitination) has been widely studied in yeast and mammalian systems. Defects in Pex1, Pex6 or Pex15/PEX26 (exportomer components) disrupt recycling of Pex5, and this results in an accumulation of the ubiquitinated Pex5 in the peroxisomal membrane in budding yeast (Platta et al., 2004; Kiel et al., 2005), while in *Pichia Pastoris* and human PBD patient cells, there is striking reduction in the steady state levels of Pex5 (Dodt et al., 1996; Collins et al., 2000). The stuck and ubiquitinated receptor is released by a quality control pathway called RADAR (receptor accumulation and degradation in the absence of recycling) (Leon et al., 2006). The effect of PEX1 or PEX6 depletion on the steady state levels of PEX7 and the role of the RADAR pathway for quality control of PEX7 remained poorly characterized. The PTS2-import receptor PEX7 requires a co-receptor in all organisms, which is Pex18p/Pex21p (*S. cerevisiae*), PEX20 (*Y. lipolytica* and other yeasts) (Einwachter et al., 2001) or PEX5 itself in humans (Braverman et al., 1998) and plants (Woodward et al., 2005). In humans, two isoforms of PEX5 are produced by alternative splicing, i.e., PEX5L and PEX5S. Only the long isoform HsPEX5L contains a PEX7 binding box. Similar to plants, trypanosomatid parasites encode a single PEX5 protein, which also harbors a PEX7-binding motif (Galland et al., 2007). PEX5 is shown to be ubiquitinated in trypanosomes (Gualdron-Lopez et al., 2013). Regarding PTS2 pathway, co-receptors PEX18/PEX20 are known to be ubiquitinated (Hensel et al., 2011; El Magraoui et al., 2013; Liu et al., 2013). It can be envisaged that, in the absence of PEX1 in trypanosomes, PEX5 and PEX7 remains stuck at the glycosomal membrane and gets poly-ubiquitinated. This would further signal for the recruitment of a quality control machinery that dislocates PEX5 along with PEX7 from the glycosomal membrane, further destined for proteasomal degradation. Accordingly, our study shows that both import receptors steady state levels fall below detection limit upon PEX1 RNAi knockdown. Our data show that also trypanosomes contain the RADAR pathway, which takes care of the removal of the import receptors that are stuck at the glycosomal membrane. The data also show that the RADAR pathway, at least in trypanosomes, is not only responsible for quality control of the PTS1 receptor PEX5 but also for the PTS2-receptor PEX7. For PEX5, it is known that the released receptor is degraded by the proteasome (Platta et al, 2004; Kiel et al, 2005; Leon et al, 2006). However, it is still unknown how the receptor is released from the membrane. A possible candidate could be ATAD1/Msp1 (Grimm et al., 2016; Weidberg et al., 2018) ortholog, which is so far not characterized in the trypanosomatid parasites.

*Tb*PEX6 and *Tb*PEX1 do not contain predicted transmembrane domains, yet they localize to the glycosomes. This suggests that they are anchored to the glycosomal membrane by an adaptor protein. In other organisms, this anchoring is performed by Tail-anchored (TA) proteins Pex15p/PEX26/APEM9 (Birschmann et al., 2003; Matsumoto et al., 2003; Goto et al., 2011). For yeast, it has been shown that Pex1p and Pex6p form a stable complex in the cytosol, and this complex binds to the peroxisomal membrane anchor Pex15p (Rosenkranz et al., 2006). Furthermore, Pex15p also associates with the components of importomer, thus bridging it with the exportomer. The similarity of the trypanosomal PEX1 and PEX6 to its counterparts in other species allow to predict the existence of such a membrane anchor also in parasites, however, primary sequence as well as structure-based homology searches so far failed to identify *Trypanosoma* counterpart. Regarding the druggability against trypanosomatid parasites, the PEX1 ATPase activity or the PEX1-PEX6 interaction might be poor drug targets, due to high degree of conservation with the human counterpart. However, the interaction of PEX1-PEX6 complex with the glycosomal membrane anchor could be an attractive target, as it appears to be highly divergent.

## Author contributions

LM, VK and RE conceived and planned the experiments. LM performed all experiments, HA performed the bioinformatic analysis, WS, RE VK supervised the work. LM and VK wrote the manuscript with support from WS and RE. All authors read and gave feedback on the manuscript.

## Acknowledgements

We thank Prof. Paul Michels for kindly providing various antibodies and Nadine Schmidt for providing assistance in ELYRA microscopy.

