## Supplementary Material for "PEX1 is essential for the glycosome biogenesis and trypanosomatid parasite survival"

#### Supplementary Figures:

**A.**

| Organism | *T. brucei* | *T. cruzi* | *L. donovani* | *H. sapiens* | *S. cerevisiae* | *A. thaliana* | *D. melanogaster* |
|---|---|---|---|---|---|---|---|
| ***T. brucei*** | | 23.7% | 16.0% | 13.8% | 13.3% | 16.1% | 13.0% |
| ***T. cruzi*** | 17.1% | | 15.7% | 12.6% | 14.5% | 14.5% | 15.0% |
| ***L. donovani*** | 9.4% | 9.4% | | 15.5% | 14.4% | 16.3% | 14.6% |
| ***H. sapiens*** | 5.6% | 7% | 8.1% | | 16.4% | 15.9% | 12.6% |
| ***S. cerevisiae*** | 5.9% | 5.7% | 6.7% | 7.5% | | 14.3% | 14.5% |
| ***A. thaliana*** | 6.8% | 7.2% | 7.8% | 6.9% | 5.9% | | 14.1% |
| ***D. melanogaster*** | 5.6% | 5.9% | 7.2% | 4.8% | 5.5% | 5.8% | |

**% IDENTITY** (lower triangle)

**% SIMILARITY** (upper triangle)

**B.**

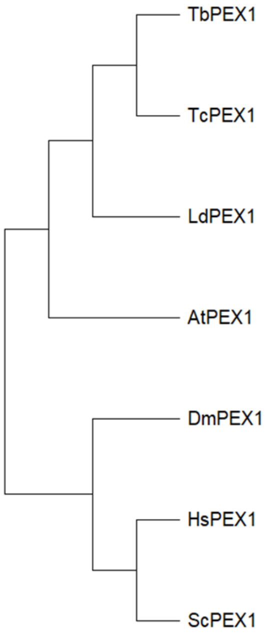


**Supplementary Figure 1. A.** PEX1 sequence identity and similarity matrix based on SIAS homology modelling **B.** Phylogenetic tree construction of the putative PEX1 proteins from *Trypanosoma brucei (Tb)*, *Trypanosoma cruzi (Tc)*, *Leishmania donovani (Ld)* and known PEX1 proteins from *Saccharomyces cerevisiae (Sc)*, *Homo sapiens (Hs)*, *Arabidopsis thaliana (At)*, and *Drosophila melanogaster (Dm)*.

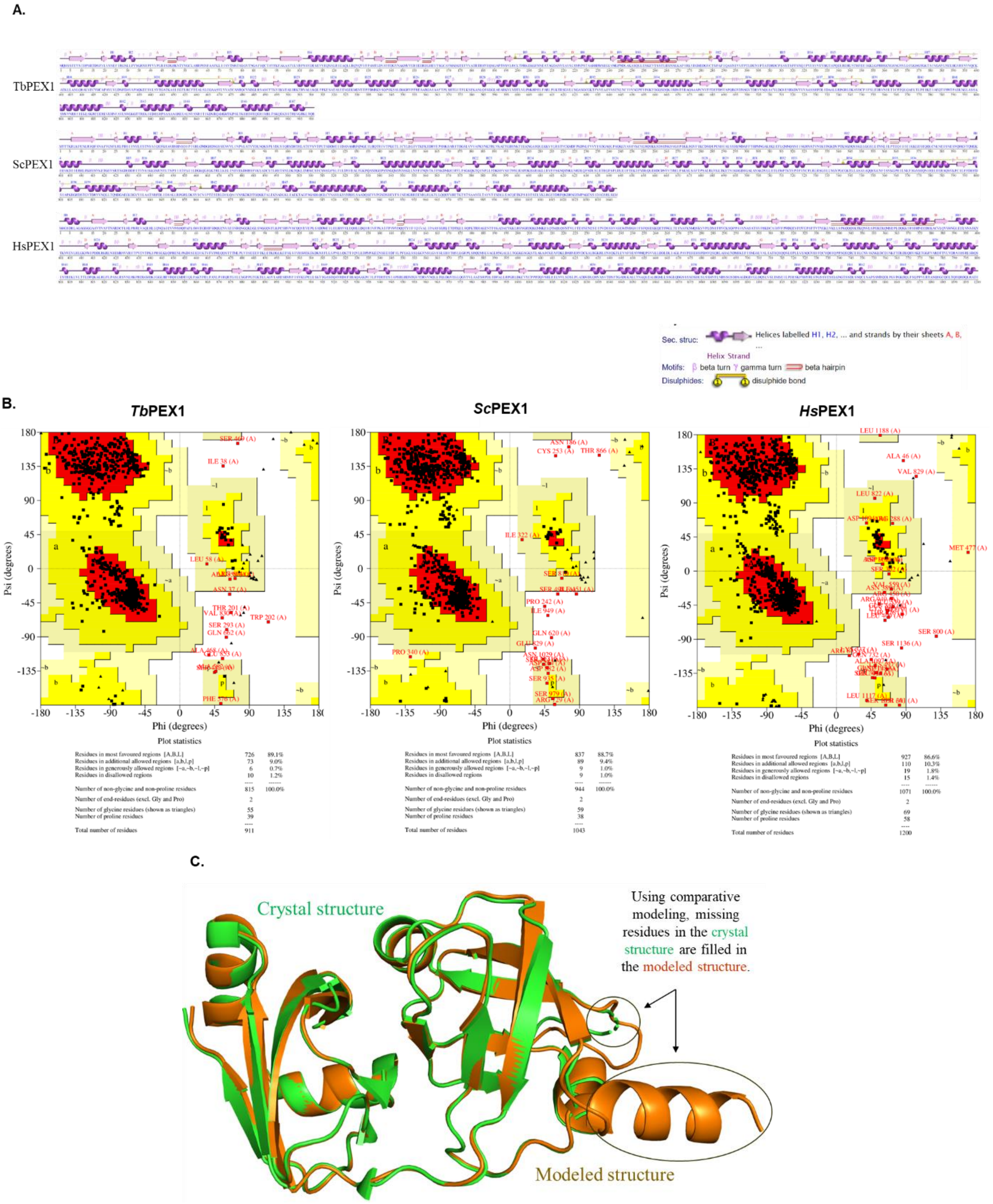


**Supplementary Figure 2. A.** 2D structure prediction of the putative PEX1 protein of *Trypanosoma brucei*, and the known PEX1 proteins of *Saccharomyces cerevisiae* and *Homo sapiens* **B.** Ramachandran plot for modelled PEX1 protein of *Trypanosoma brucei*, *Saccharomyces cerevisiae* and *Homo sapiens* obtained from Saves server of PROCHECK validation package. Less than 1.5% of residues were present in the Ramachandran plot's disallowed region **C.** Comparative modelling of the N-terminus of PEX1 3D structure showing the superimposition of the crystal structure of mouse PEX1 NTD (green) with the modelled structure of same protein (brown) with RMSD value of 0.29 Å. Same superimposition was also seen for the modelled structure of *Hs*PEX1 (95% sequence identity with mouse PEX1 NTD) with the crystal structure of mouse PEX1 NTD.

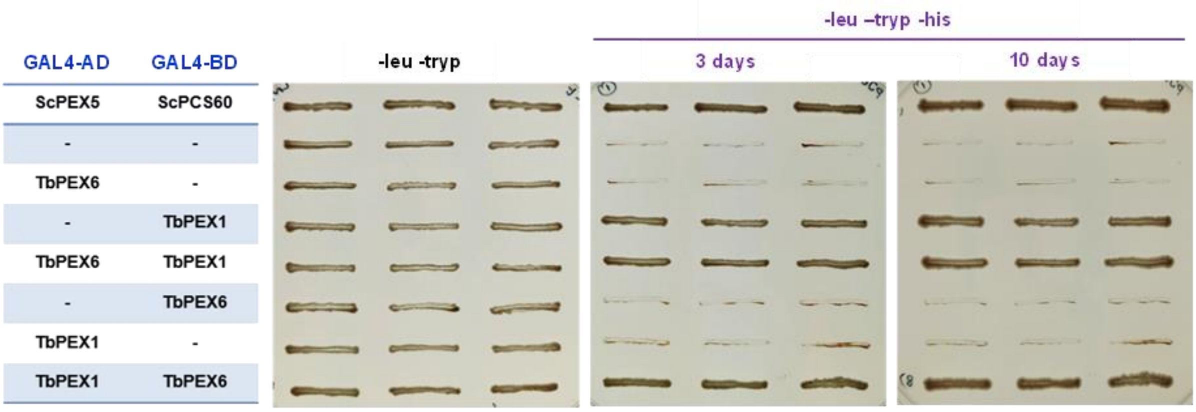


**Supplementary Figure 3. Interaction of *Tb*PEX1-*Tb*PEX6 in the Yeast Two-Hybrid system.** PJ694a (yeast strain) double transformed clones were grown in -leu -tryp plates and in -leu -tryp -his dropout plates containing 3-amino-1,2,4 triazole. Cell growth after 3 days and 10 days indicates the interaction between *Tb*PEX6 (AD) and *Tb*PEX1 (BD). The interaction between ScPEX5 and its cargo ScPCS60 is used as a positive control.

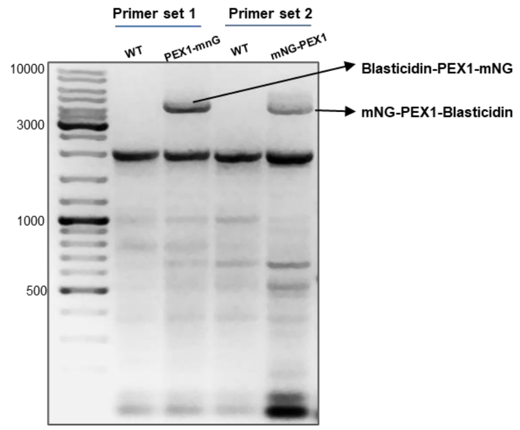


**Supplementary Figure 4.** PCR confirmation of endogenous tagging of *Tb*PEX1 with mNeonGreen in BSF trypanosomes using Primer set 1 (RE 7224-RE7225) and Primer set 2 (RE 7226-7227).

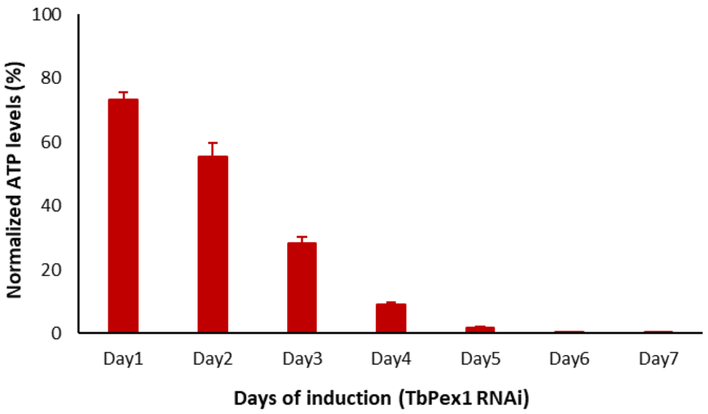


**Supplementary Figure 5.** Cell viability assessment of cells induced for PEX1 RNAi by CellTitre-Glo method. The viability measurement of RNAi induced cells (Tetracycline addition) were normalized to non-induced cells (DMSO treatment) in the respective days (day1 to day 7). Error bars represent the Standard deviation of the three biological replicates.

### Supplementary Tables:

#### Table 1 Primers

| No. | Primer name | Sequence (5'→3') |
|---|---|---|
| 1 | RE7071 | ACGATGTCGACAATGCAGCACAGCTCCTTTGAA |
| 2 | RE7072 | ATAATGAGCTCTCACCGTTGGGTCAATTTCCTTCC |
| 3 | RE7073 | ATCGAGTCGACCATGATGTCACGGACCGTTGAG |
| 4 | RE7074 | ATAATACTAGTTCAATCAGCCACACGACCCGA |
| 5 | RE7224 | AACAGGAAAGAATTCACCGCCTCTTCAGTAAGCAGGAAAAGAGCAGTACGAGGGA AGTCGGAAGGAAATTGACCCAACGGGGTTCTGGTAGTGGTTCC |
| 6 | RE7225 | GAGCCCCAAGCCATACTGGAGAGGGCAGAGGAGTGGCTGACAGAAATTAACGAAT TGTAGAGTGTTTTGGTGTTACCACCCCAATTTGAGAGACCTGTGC |
| 7 | RE7226 | TTTCTTCGTTCTCCAACCGATCTAGAGGGTCCCTTTATAAACGTAATTATTACTAGT GTAGCTCCAAAAAGAATAAGAGGGTATAATGCAGACCTGCTGC |
| 8 | RE7227 | CGGATGAATTCATTTGACACCAAGACAAATGAGTCCGTTCGGGACGGGTCGATAGA AACTTCAAAGGAGCTGTGCTGCATACTACCCGATCCTGATCC |
| 9 | RE7323 | ATGACAAAGCTTCATTGGCCAGAGTGAGCAGA |
| 10 | RE7324 | ATGACAGGGCCCCCATACTCGCTGATGACGCT |
| 11 | RE7325 | ATGACAGGATCCCATTGGCCAGAGTGAGCAGA |
| 12 | RE7326 | ATGACAGGGCCCACGCTCCTCACAACGCTTGGA |

#### Table 2 Strains and Plasmids

| No. | Expression system | Construct | Primers | Restriction sites | Vector used for cloning |
|---|---|---|---|---|---|
| 1 | *Saccharomyces cerevisiae* | GAL4AD-*Tb*PEX1 | RE7071-RE7072 | SalI/SacI | PC86 |
| 2 | *Saccharomyces cerevisiae* | GAL4BD-*Tb*PEX1 | RE7071-RE7072 | SalI/SacI | PC97 |
| 3 | *Saccharomyces cerevisiae* | GAL4AD-*Tb*PEX6 | RE7073-RE7074 | SalI/SpeI | PC86 |
| 4 | *Saccharomyces cerevisiae* | GAL4BD-*Tb*PEX6 | RE7073-RE7074 | SalI/SpeI | PC97 |
| 5 | *Trypanosoma brucei* | *Tb*PEX1-mNG | RE7224-RE7225 | - | pPOTV7 |
| 6 | *Trypanosoma brucei* | mNG-*Tb*PEX1 | RE7226-RE7227 | - | pPOTV7 |
| 7 | *Trypanosoma brucei* | Fragment I (*Tb*PEX1 Stem loop RNAi) | RE7323-RE7324 | HindIII/ApaI | pHD1336 |
| 8 | *Trypanosoma brucei* | Fragment II (*Tb*PEX1 Stem loop RNAi) | RE7325-RE7326 | ApaI/BamHI | pHD1336 |
